# Shipment Stress in Early Life Aggravates Disease Pathogenesis in Mice with Experimental Autoimmune Encephalomyelitis: Support for a Two-Hit Hypothesis of Multiple Sclerosis Etiology

**DOI:** 10.1101/2023.05.22.541749

**Authors:** Jamshid Faraji, Dennis Bettenson, V. Wee Yong, Gerlinde A. S. Metz

**Author notes:** **Corresponding authors: Gerlinde A.S. Metz**, PhD, Canadian Centre for Behavioural Neuroscience, University of Lethbridge, 4401 University Drive, Lethbridge, Alberta T1K 3M4, Canada, **Jamshid Faraji**, PhD, Canadian Centre for Behavioural Neuroscience University of Lethbridge, 4401 University Drive, Lethbridge, Alberta T1K 3M4, Canada.

## Abstract

Visual impairments are one of the earliest diagnosed symptoms of multiple sclerosis (MS). The onset and progression of vision loss in MS may be influenced by cumulative psychophysiological stress. Here, we used a two-hit model of stress in female mice to determine if early life stress (ELS) influences the clinical severity of experimental autoimmune encephalomyelitis (EAE) later in life. We hypothesized that ELS caused by animal transportation during early postnatal development represents a co-factor which can exacerbate the disease severity of EAE. Adult EAE mice with ELS displayed more severe clinical signs and delayed recovery compared to non-stressed EAE mice. ELS also diminished visual acuity measured by optokinetic responses, locomotion and exploratory behaviours in EAE mice. Notably, ELS caused earlier onset of visual impairments in EAE. Exacerbated functional impairments in stressed EAE mice were highly correlated with circulating corticosterone levels. The findings show that the progression of induced EAE (second hit) in adulthood can be significantly impacted by adverse early life experiences (first hit). The observations emphasize the importance of comprehensive behavioural testing, including non-motor functions, to enhance the translational value of preclinical animal models of MS. Moreover, shipment stress of laboratory animals should be considered a necessary variable in preclinical MS research. The consideration of cumulative lifetime stresses provides a new perspective of MS pathogenesis within a personalized medicine framework.

## Introduction

Multiple sclerosis (MS) is a chronic immune-mediated demyelinating disease which is characterized by progressive neurodegeneration and neurological disability. One of the earliest symptoms of MS is optic neuritis, which often results in significant vision loss (Redler & Levy, 2020). Although many pathological processes have been characterized, the main causes of MS are still not fully understood (Reich et al., 2018). Aside from genetic predisposition, the main contributors to autoimmune diseases arguably involve environmental and lifestyle factors, such as adverse life experiences (Heynen et al., 2022; Verstraeten et al., 2019). For example, exposure to early life stress (ELS) increases the vulnerability to psychiatric (Goodwill et al., 2019; Peña et al., 2019; Zhang et al., 2016) and neurological diseases (Faraji et al., 2022b; Malter Cohen et al., 2013; Paladini et al., 2022; Peña et al., 2017). ELS can alter hypothalamic-pituitary-adrenal (HPA) axis function (Faraji et al., 2017a) and sensitize specific neurocircuits to subsequent stress (Ladd et al., 2005), and even change transcriptomic patterns in the brain (Peña et al., 2019). All of these changes can potentially alter immune responses and the risk of autoimmune diseases. Further, early life stress may challenge immune function later in life by changing inflammatory responses (Danese & S, 2017). Such findings offer a framework through which inflammatory response patterns may predict the pathophysiology and neuroimmunological symptoms of MS (Faraji et al., 2022a; Kuhlmann et al., 2023).

ELS and repeated stress later in life may threaten homeostatic regulation of immune responses and eventually initiate pathological processes, according to the concept of allostatic load (Karatsoreos & McEwen, 2011). Thus, ELS may augment HPA axis responses when combined with a second stressor, such as neuronal damage or a pro-inflammatory event, in adulthood (Faraji et al., 2017b). Our work has shown that the effects of prenatal stress can remain silent across four generations but generate a phenotype that is vulnerable to psychopathology in adulthood when challenged with a second stressor (Faraji et al., 2017a). Hence, the interaction between two hits of stress, such as a psychophysiological stressor and an immune challenge, may increase vulnerability to physiopathology and psychopathology while individually create minimal systemic challenges (Verstraeten et al., 2019). According to the two-hit hypothesis (Cotella et al., 2014; Faraji et al., 2017b; Giovanoli et al., 2013), therefore, latent susceptibility to physiological stress in later life may be intensified by ELS. Our previous observation (Gerrard et al., 2017) showed that postnatal stress synergistically exacerbates the severity of experimental MS in rats via altering epigenetic regulatory pathways suggesting that stress can represent a significant risk factor for symptomatic deterioration in clinical MS.

Although females are 2-3 times more susceptible to MS than males (Doss et al., 2021), much of the previous research has focused on males. The heightened vulnerability among females to MS has been also reported in the experimental autoimmune encephalomyelitis (EAE) model, a common preclinical model of human MS (Wiedrick et al., 2021). In the present study, we investigated whether ELS induced by psychophysiological stress in female mice can diminish recovery in a mild form of EAE. We propose that psychophysiological and immune stresses present a two-hit model for symptomatic EAE with the following components: (*i*) an early-life psychophysiological challenge induced by shipment stress (the first hit), and (*ii*) an immune insult induced by EAE (the second hit) later in life. We hypothesized that ELS leads to greater clinical severity of EAE in adulthood and HPA-axis hyperactivity in female mice. The findings provide the first evidence that adverse early life experiences can exacerbate the consequences of a neuroimmunological insult later in life, proposing a mechanism for risk susceptibility to MS in adulthood.

## Results

### EAE-Induced Stress and Immune Responses are Exacerbated by Early Life Stress

Figure 1 illustrates CORT levels in all groups across different time points (pre-induction, day 19 and day 27) and spleen weight. Six mice in different groups were excluded from the blood sampling and CORT analysis due to technical issues. A Kruskal-Wallis H test showed no significant differences between groups in pre-induction (baseline) CORT (*p*=0.394; *panel A*). However, there was a statistically significant group difference (*H*_3_=14.89, *p*<0.01) with a mean rank CORT level of 5.20 for No Stress, 18.83 for No Stress+EAE, 8.83 for Stress and 19.45 for Stress+EAE on day 19 post-induction. Multiple comparisons also showed that EAE groups were different from No Stress groups (all p≤0.01; *panel B*). Also, there was a significant difference between groups on day 27 (*H*_3_=11.29, *p*<0.01) where mean rank CORT levels were 8.60 for No Stress, 14.00 for No Stress+EAE, 6.20 for Stress and 18.56 for Stress+EAE groups. Multiple comparisons showed a significant difference between Stress and Stress+EAE groups (*p*<0.01; *panel C*).

**Figure 1:**
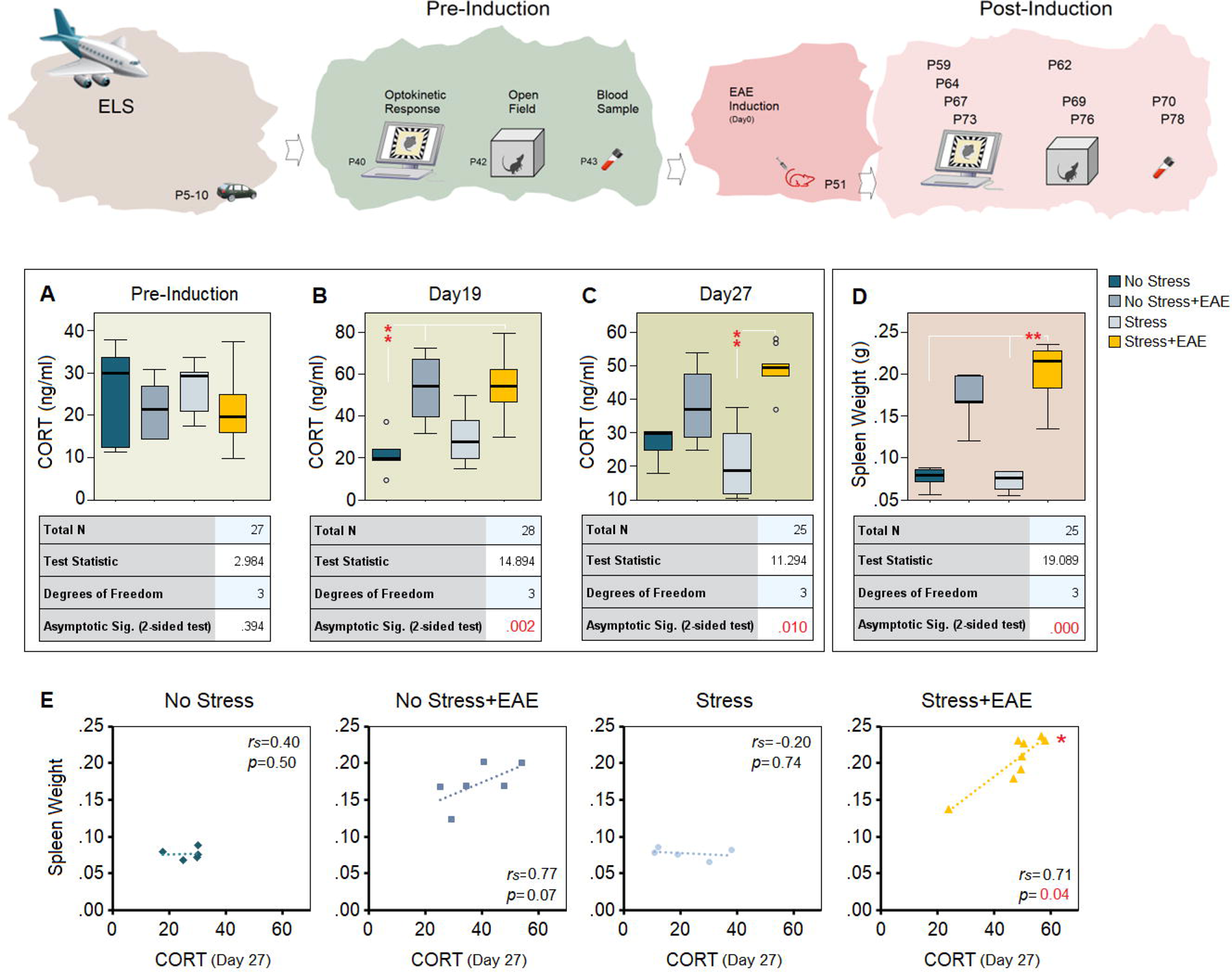
Markers of stress and immunological activity before and after EAE induction. (A-C) Both No Stress+EAE mice (*n*=9) and Stress+EAE mice (*n*=8) displayed a strong CORT response to EAE, particularly on day 27 post-induction. (D) Spleen weight increased in Stress+EAE, mice only. (E) Correlational analysis shows that splenomegaly in response to early life stress and EAE can be predicted by HPA hyperactivity and over-production of CORT on day 27 post-induction. Asterisks indicate significant differences: \**p*≤0.05, \*\**p*≤0.01; *Kruskal-Wallis H test* and *Spearman’s rank correlation*.

Further, spleen weight revealed a significant difference between groups (*H*_3_=19.08, *p*<0.001) with a mean rank spleen weight of 6.80 for No Stress, 16.00 for No Stress+EAE, 5.33 for Stress and 20.38 for Stress+EAE groups. Again, multiple comparisons indicated significant differences between Stress+EAE, and Stress and No Stress groups (all p≤0.01; Figure 1 *panel D*). Thus, Stress+EAE mice responded to the EAE challenge with higher levels of CORT and greater splenic weight than other groups. Also, Spearman’s rank correlation analysis revealed a significant relationship between spleen weight and CORT levels on day 27 post-induction in Stress+EAE group (*r*s=0.71, *p*≤0.05; Figure 1 *panel E*) indicating that HPA chronic hyperactivity in response to ELS significantly predicts splenomegaly in EAE. The enlarged spleen may indicate inflammatory deficiency influenced by HPA axis activation in Stress+EAE mice.

### Early Life Stress Reduces Recovery Following EAE

EAE disease progression is shown in Figure 2*A*. The peak of clinical disabilities occurred between days 15–17 post-induction. At the peak of disability, animals in Stress+EAE and No Stress+EAE groups showed partially to fully flaccid tails, locomotor difficulty and paralysis in at least one hind limb. However, Stress+EAE mice (*n*=11) displayed a tendency to maintain higher disease scores than No Stress+EAE mice (*n*=8) until day 22 post-induction. Disease scores indicated a significant group difference on days 21 to 25 when Stress+EAE mice displayed more severe deficits (higher scores) than No Stress+EAE animals (*U*=17.50, *U*=19, *U*=19.50, *U*=19.50, *U*=19.50, all *p*≤0.05; Mann-Whitney U). The CDI on post-induction days 7–26 indicated more severe symptoms in Stress+EAE than No Stress+EAE mice (mean rank: 11.18 *vs.* 8.38), even though the observed difference was not statistically significant (*U*=31, Z=-1.07, *p*≥0.28; Mann-Whitney U, Figure 2 *panel B*). All animals without EAE remained asymptomatic with 0 clinical scores (data not shown).

**Figure 2:**
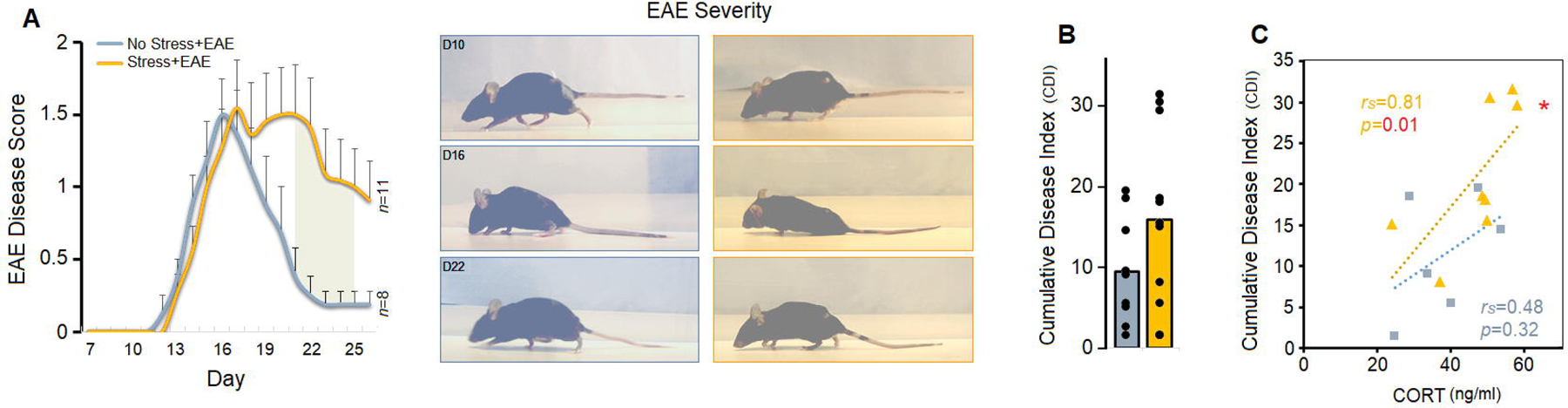
Disease progression of EAE in stressed and non-stressed mice. (A) Stress+EAE animals (*n*=11) displayed significantly higher disease scores than No Stress+EAE mice (*n*=8) on days 21-25 post-induction (grey area). Right panel compares EAE severity in a representative mouse from each group on three post-induction days (days, 10, 16, and 22). *Day 10*: the tail was mobile, and no tail paralysis was observed. *Day 16*: partial or complete rigidity was felt in the tail in both groups. However, limp tail and hind limb paralysis was more pronounced in Stress+EAE mice. In most stressed animals, the hind limbs were not weight bearing, and tail movement was absent. *Day 22*: in both groups, forelimbs and hind limbs were used for ambulation, and animals were able to regain weight support with hind limbs. However, the tail was still limp in the stressed animals. (B) Stress+EAE mice obtained higher CDI scores. Black circles represent individual animals. EAE in stressed animals revealed a significantly accelerated progression and aggravated symptom severity. (C) The greater CDI in Stress+EAE mice was significantly associated with increased levels of CORT on day 27 post-induction. Asterisks indicate significant differences: \**p*≤0.05; *Mann-Whitney U test* and *Spearman’s rank correlation*.

Correlations between EAE severity and HPA axis activity revealed a significant positive relationship between CDI and CORT levels on day 27 in Stress+EAE mice (*r*s=0.81, *p*≤0.01; Spearman’s rho; Figure 2 *panel C*) suggesting that greater CDI in the Stress+EAE animals was associated with higher levels of CORT at the chronic time point post-induction (day 27). It appears that EAE in stressed mice followed the characteristic symptomatic progression with slower recovery process (delayed remission) influenced by ELS exposure.

### Early Life Stress Predicts Earlier and Prolonged Vision Impairments in EAE Mice

Figure 3*A-D* show that visual acuity gradually declined in both EAE groups, and in both groups the two acuities (CW and CCW) were not identical. *R-M* ANOVA indicated a main effect of Group (*F*_1,19_=236.63, *p<*0.001, η^2^=0.92) and Time Point (*F*_3,57_=7.57, *p<*0.01, η^2^=0.57), but no interaction between Group and Time Point (*p<*0.24). The differences observed between the directions of motion (CW *vs*. CCW) also were not significant. A separate *O-W* ANOVA conducted to compare frequency threshold on post-induction days 8 (*p<*0.01), 13 (*p<*0.001), 16 (*p<*0.01), and 22 (*p<*0.05) found that mice with ELS had lower frequency thresholds across all post-induction time points (*panel B)*. Thus, ELS postponed improvement in visual function at post-induction days when compared with non-stressed animals. Also, visual acuity was negatively correlated with CDI such that animals in both groups with higher visual acuity with CW motion would have lower CDI (No Stress+EAE: (*r*s =-0.90, *p*≤0.05, Stress+EAE: *r*s=-0.76, *p*≤0.05, Spearman’s rho; *panel C*). Alternatively, the frequency threshold when the direction of motion was set on CW significantly predicted chronic levels of CORT (day 27) only in No Stress+EAE mice (*r*s=-0.94, *p*≤0.01; Spearman’s rho) suggesting that non-stressed animals with higher frequency threshold displayed HPA axis hyperactivity. Such positive relationship profile was also observed in stressed animals with EAE in both directions of motion, even though the correlation was not significant (*panel D*).

**Figure 3:**
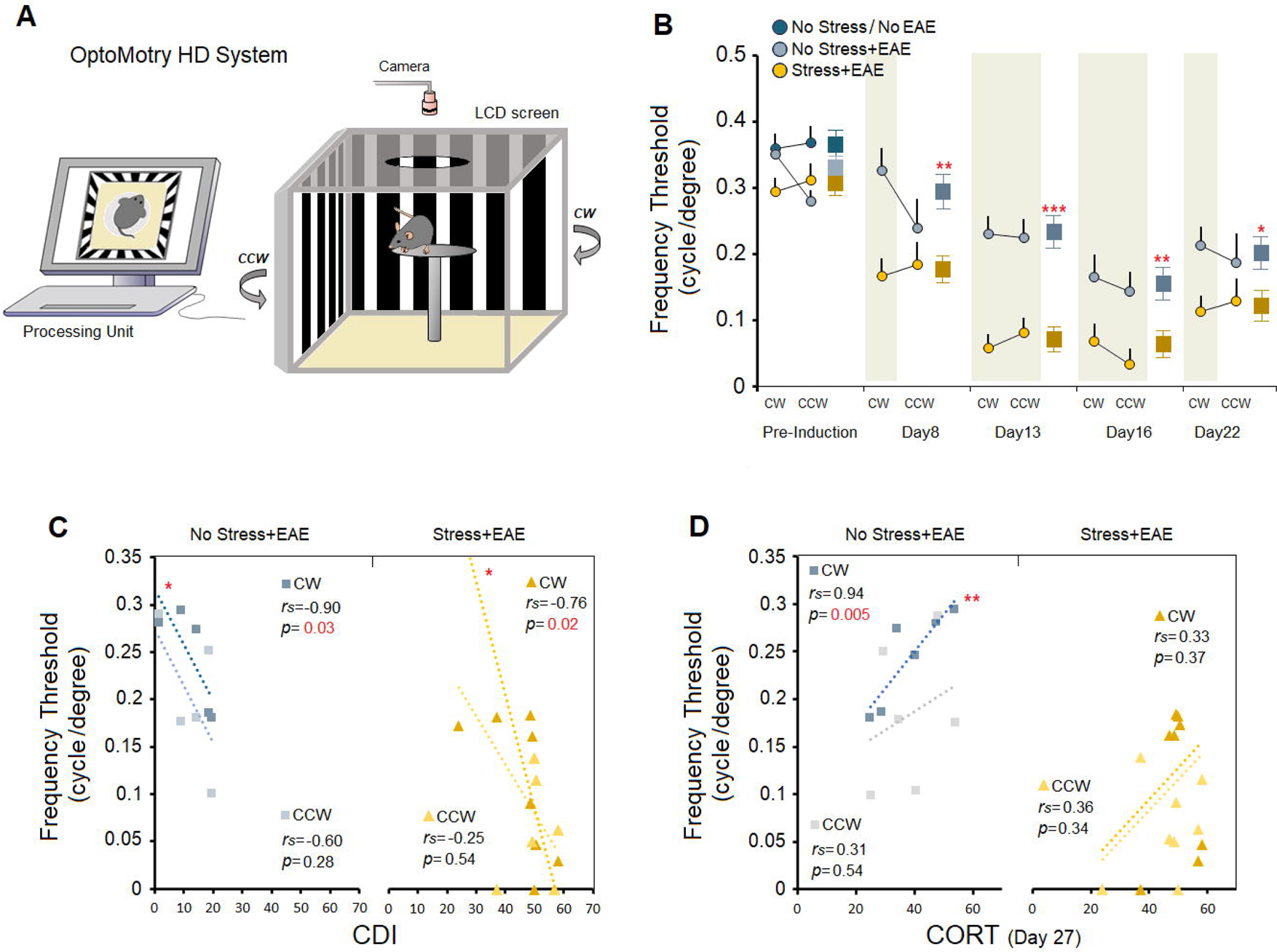
Visual acuity and optokinetic head tracking. (A) Configuration (top and side view) of the virtual optomotor system with a visual stimulus unit which displays rotating black-and-white strips clockwise (CW) and counterclockwise (CCW) on four LCD screens. (B) EAE induction reduced the frequency threshold (visual acuity) in both groups; however, Stress+EAE mice (*n*=13) displayed significantly decreased visual acuity at all time points post-induction. Average frequency threshold is also shown at each time point. Grey boxes represent statistically significant differences between groups. Panels C&D show the correlations between frequency threshold and CDI, and frequency threshold and CORT, respectively. Asterisks indicate significant differences: \**p*≤0.05, \*\**p*≤0.01, \*\*\**p*≤0.001; *O-W* and *R-M* ANOVA, and *Spearman’s rank correlation*.

### Early Life Stress Reduces Locomotor Activity and Recovery Following EAE

Locomotor activity and exploratory behaviour were assessed by overall activity or total distance traveled within an open field. *Repeated measure* ANOVA did not show a significant main effect of Group, but Time Point (*F*_3,54_=54.35, *p<*0.001, η^2^=0.75) and an interaction between Group and Time Point (*F*_3,54_=4.45, *p<*0.01, η^2^=0.19). However, Stress+EAE mice (*n*=11) traveled significantly less than No Stress+EAE mice (*n*=9) at the chronic time point post-induction (day 25; 494.69 ± 82.11 cm *vs.* 737.77 ± 67.69 cm, *F*_1,20_=4.54, *p<*0.05, *O*ne *Way* ANOVA; Figure 4*A*). Also, rate of change (ROC) during all three time points after EAE induction showed that Stress+EAE mice experienced greater reduction in exploratory behaviours from pre-induction to post-induction day 25. The ROC analysis also indicated that, unlike No Stress+EAE mice, post-induction recovery in exploratory behaviour was reduced in Stress+EAE mice. The No Stress+EAE mice displayed a profile of recovery to approximately 32% on day 25 relative to day 18, whereas the Stress+EAE animals showed only 6% recovery at the same time (Figure 4 *panel B*). Moreover, Spearman’s rho correlation coefficient showed that increased plasma CORT reliably predicted increased locomotion in Stress+EAE and No Stress+EAE groups, however, the observed correlation was significant exclusively in the No Stress+EAE group (*r*s=0.82, *p*≤0.04; Figure 4 *panel C*).

**Figure 4:**
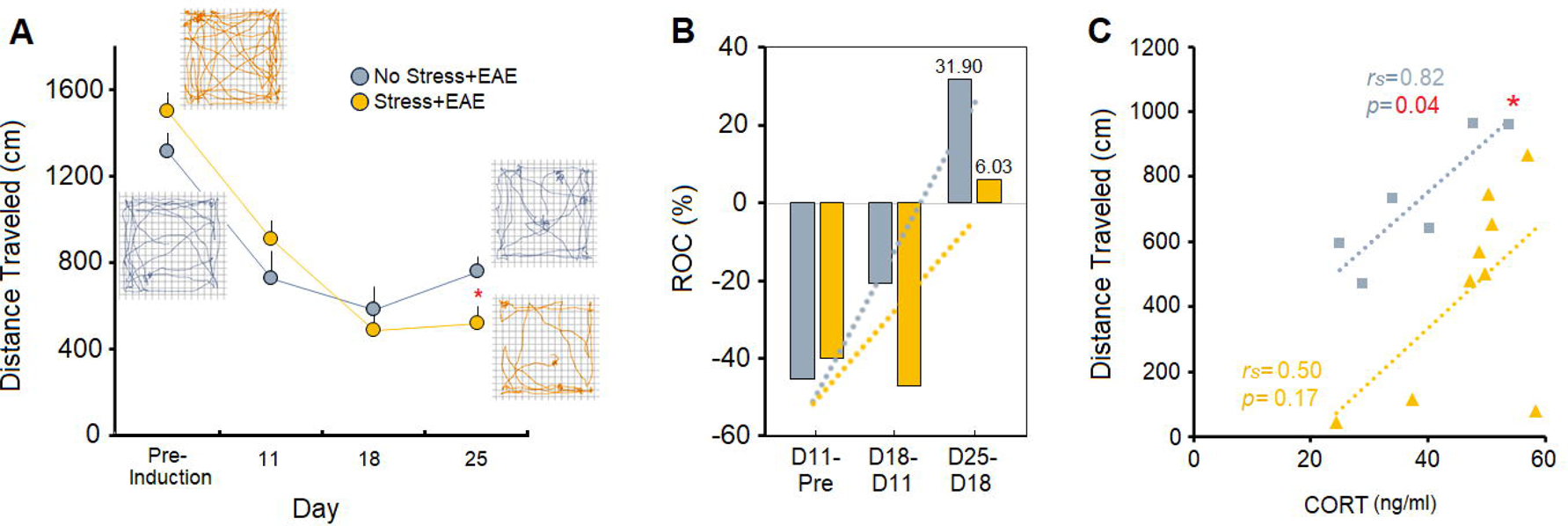
Locomotor activity and exploration before and after EAE induction. (A) Analysis of exploratory behaviours indicated that path length in the open field was impacted by EAE at all post-induction time points. Stress+EAE mice (*n*=11), however, traveled significantly shorter distances than No Stress+EAE mice (*n*=9) on post-induction day 25. The open field exploration paths shown in the motion track graphics compare the trajectory made by one animal in each group before EAE induction versus post-induction day 25. (B) The rate of changes (ROC) on each post-induction day relative to the previous time point (day 11 to pre-induction, day 18 to day 11, and day 25 relative to day 18) shows that ELS exposure limited the recovery from EAE on post-induction day 25 in Stress+EAE mice. (C) The distance traveled by both groups was positively correlated with plasma CORT, however, changes in exploratory behaviours after EAE induction only predicted the chronic CORT levels in No Stress+EAE mice. Asterisks indicate significant differences: \**p*≤0.05; *O-W* ANOVA and *Spearman’s rank correlation*.

## Discussion

Evidence is lacking for the cumulative impact of early postnatal adversities and neuroimmunological challenges in adulthood, such as MS. In the present study, we showed that early life shipment stress in female mice leads to aggravated risk and diminished recovery following EAE in later life. Notably, ELS promotes earlier onset of vision impairments, one of the earliest symptoms of MS in patients. Mice with ELS also displayed higher susceptibility to the clinical course and severity of a mild form of EAE compared to non-stressed mice (*Fig.5*). The differential effect among groups suggests that even a moderate stress during the early postnatal days (first hit) promotes a vulnerable phenotype that is more susceptible to an immunological challenge (second hit), followed by exacerbated deficits and reduced recovery in both sensory and motor abilities. The heightened susceptibility to EAE in stressed mice was also reflected in their greater level of CORT responses to the disease during the chronic phase. Our findings reveal synergistic interactions between two stressors with cumulative impact on EAE severity, and support theories that onset and progression of pathogenesis in many diseases involve multiple or accumulated psychophysiological hits (Giovanoli et al., 2013; Guerrin et al., 2021; Lopes et al., 2022; Olson et al., 2015; Picci & Scherf, 2015).

**Figure 5:**
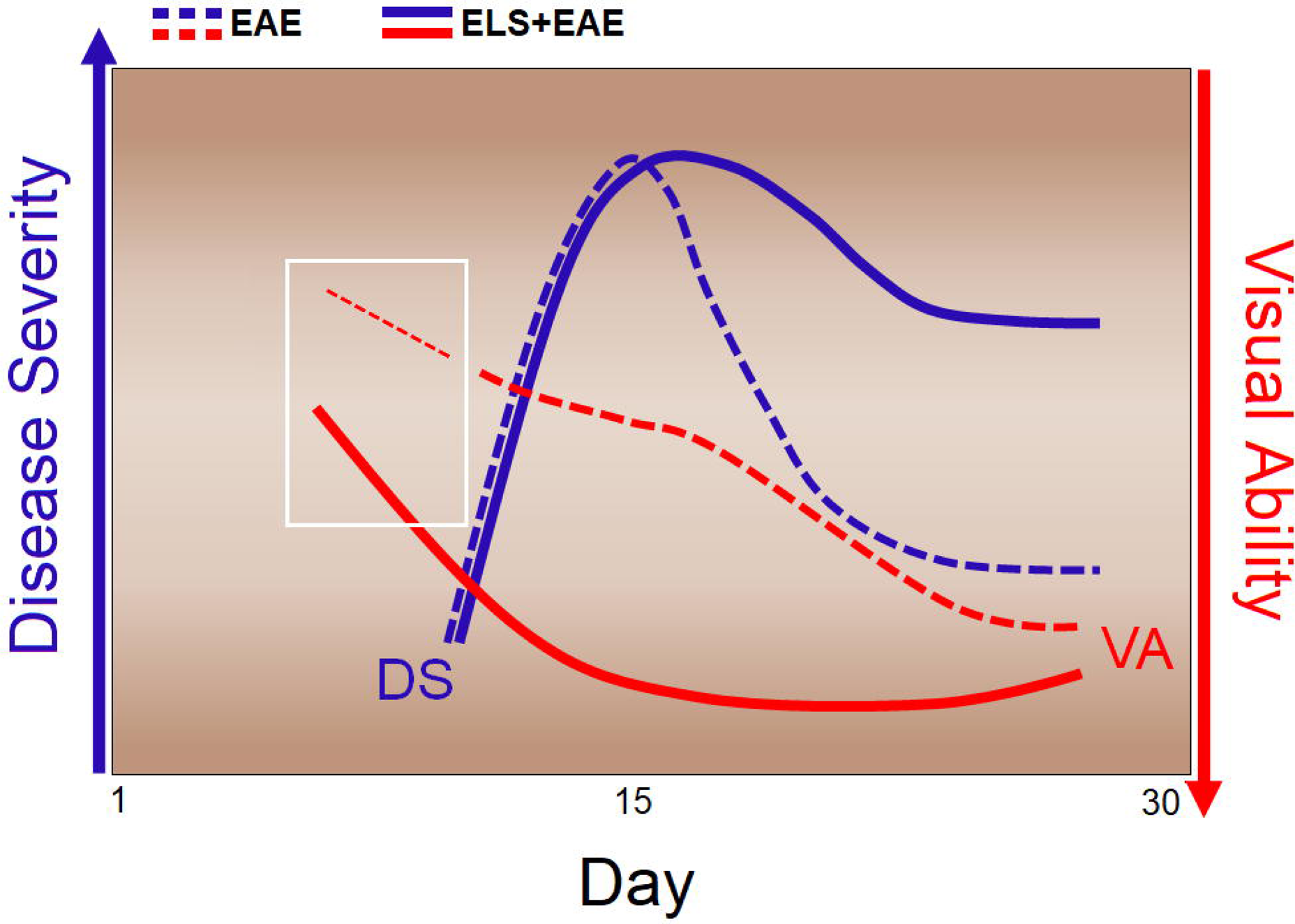
Representation of findings in the present study. Adult EAE mice with ELS displayed more severe clinical signs and delayed recovery compared to non-stressed EAE mice. ELS also promotes earlier onset of vision impairments, one of the earliest symptoms of MS in patients.

A major source of disability that is commonly seen with a female preponderance ratio in MS patients is optic neuritis (Ciapă et al., 2022; Redler & Levy, 2020; Sarrazin et al., 2022). Optic neuritis is a demyelinating disorder characterized by inflammation of the optic nerve, and clinical presentation of vision defects such as unilateral loss of visual acuity, visual field and color vision deficits, and decreased contrast (Redler & Levy, 2020). Similar profiles of visual dysfunction induced by demyelination and axonal damage of the optic nerves have been also reported in EAE models (Hecker et al., 2020; Itoh et al., 2018; Joly et al., 2022; Khan et al., 2014). Using the virtual optomotor system (VOS) (Prusky et al., 2004) provides a quantitative measure of eye movements (optokinetic nystagmus) and head movements (optomotor tracking) in rodents, thus showing the properties of the retinal efferents to subcortical structures (Douglas et al., 2005). In accord with previous findings (Joly et al., 2022; McDougald et al., 2018), EAE in the present findings was associated with reduced visual acuity. However, the loss of frequency threshold in stressed animals was recorded at about post-induction day 8 which was earlier than the appearance of motor deficits on day 12 post-induction. Also, the deleterious impact of ELS on optokinetic responses to EAE was more pronounced in both directions of rotation (*Fig. 3B*). Due to the crossed subcortical projections from the eyes in mice, clockwise (CW) movement in VOS will normally drive the tracking function through the left eye, whereas counterclockwise (CCW) rotation will stimulate the right eye. Also, optomotor tracking seen in VOS is driven by the same subcortical visual pathways as optokinetic responses (Douglas et al., 2005; Prusky et al., 2004). Consequently, the long-lasting reduced frequency threshold in the stressed mice likely represents the greater impact of EAE-induced neuroinflammatory processes on the function of these subcortical pathways and the retinal output. Thus, the present findings enable insights into the dynamic interactions between cytokine function and stress hormone in response to homeostatic challenges.

Importantly, visual acuity during the CW rotation (left-eye movement) in both groups was negatively correlated with CDI. This likely indicates an asymmetry in optic neuritis, gliosis and synaptic impairments rather than systemic inflammation (Joly et al., 2022), which agrees with the mainly unilateral visual deficits induced by optic neuritis in MS patients (Redler & Levy, 2020). However, future experiments should consider less well-known neuroanatomical aspects of visual function in rodents (e.g., retinal afferents to the accessory optic system), and hemispheric asymmetries, which can also be observed in stress responses (Ambeskovic et al., 2017) This specifically highlights the importance of accessory optic system nuclei such as the nucleus of the optic tract and the dorsal terminal nucleus which contribute to horizontal tracking in rodents (Douglas et al., 2005) and their vulnerability to the neuroinflammatory dynamics involved in EAE (Khan et al., 2014). Alternatively, the positive correlation between frequency threshold and chronic levels of CORT here (*Fig. 3D*) can be explained, at least in part, by the widespread and critical role of corticosteroids in immunity and anti-inflammatory processes, particularly their suppressive effects on optic neuritis (Dutt et al., 2010). Interestingly, chronic immunomodulation by corticosteroids may also reduce the risk of recurrent optic neuritis and retinal ganglion cell damage. Thus, the heightened levels of CORT in association with frequency threshold following EAE induction may also represent a neurohormonal response that in turn alleviates some of the harmful consequences of the neuroinflammation.

Consistent with the clinical, hormonal and optokinetic findings, the assessment of locomotion also indicated that stressed mice traveled significantly less than non-stressed animals in the chronic phase of EAE (post-induction day 25). Although EAE has the potential to decrease exploratory behaviours in females more than males at the chronic time point (Faraji et al., 2022a), the present observations provide further insights by demonstrating that pre-exposure to ELS in females will further reduce exploratory activity. Also, Stress+EAE females achieved lower post-induction recovery from EAE compared to non-stressed EAE mice, as shown by the decreased ROC (6.03 *vs.* 31.90) on post-induction day 25 relative to day 18 (*Fig. 4B*). The correlation between distance traveled and CORT levels, however, illustrates the contribution of HPA-related influences to locomotion (Faraji et al., 2014), although the neurohormonal susceptibility failed to correlate with exploratory behaviour in Stress+EAE mice. The delayed recovery time in Stress+EAE mice might be the source of the lack of correlation between CORT levels and distance traveled. It is unjustified, however, to define a stress state based on only the HPA hyperactivity or elevated plasma CORT (Faraji & Metz, 2020), even though elevated CORT levels are a prominent neurohormonal correlate of stress-induced emotionality (Metz et al., 2005; Sutanto & de Kloet, 1994).

Despite the frequent use of animal shipment procedures in preclinical studies, the impact of transportation stress is rarely considered in experimental design and data interpretation. The long-lasting molecular, metabolic and behavioural consequences of shipment stress (Poplawski et al., 2023; Poplawski et al., 2020) arguably represent significant confounding factors affecting the outcome of experimental manipulations in rodent models of human disease. In addition, early life stress and immune challenges in later life, which individually may have modest effect sizes (Hoghooghi et al., 2020; Olfe et al., 2010), may exert cumulative and potentially synergistic effects in pathological conditions. The data support the two-hit model of stress that initial exposure to an early postnatal adversity such as shipping stress in rodents can render the offspring more susceptible to the pathological consequences of a second insult such as EAE. Thus, early postnatal shipping stress and EAE in adulthood caused synergistic effects through which stressed mice suffered earlier onset of vision impairments and more severe motor disability when exposed to EAE than non-stressed animals. Notably, motor deficiency in ELS mice became evident only on post-induction days 21 onward (*Fig. 2A*). The EAE-induced motor signs in both groups started approximately at day 12 and peaked at days 15-16. However, the deficits in non-stressed mice gradually improved starting from day 17 until day 25, whereas the disease severity in the ELS animals was remained steady, particularly with a significant delay for recovery on days 21-25. Therefore, EAE potentially unmasked latent pathophysiology in neurodevelopmentally vulnerable mice with early postnatal stressful history.

CORT levels, EAE severity and splenic morphological alterations may be directly correlated (Bałkowiec-Iskra et al., 2007; He et al., 2014; Hernandez et al., 2013). Hence, changes in CORT levels in stressed EAE mice, which in itself is associated with HPA hyperactivity, may also partially explain the higher spleen weight in the present study. In accordance with our recent findings (Faraji et al., 2022a) through which female mice responded to EAE with greater adrenal gland and splenic weights than males, we also show here that female mice with ELS exposure and EAE displayed the same profile of neurohormonal and splenic abnormality during CNS inflammation. ELS was shown to promote pro-inflammatory processes (Ambeskovic et al., 2020) and exacerbate EAE symptom severity (Paladini et al., 2022). Likewise, animals who experienced ELS exhibit elevated pro-inflammatory cytokine signaling in key brain regions (Dutcher et al., 2020) that may influence the response to neurological conditions later in life. ELS also disrupts HPA-axis responsiveness in mice, and is associated with increased peripheral inflammatory cytokine secretion and cytokine levels in brain tissue in adulthood (Hohmann et al., 2017). In absence of a precise assessment of inflammatory signatures corresponding to the neuroimmunological processes, it is difficult to draw a solid conclusion about the inflammation-related pathogenesis in ELS mice. However, one can speculate that the splenomegaly in the present experiment might not only be linked to the overproduction of CORT, but also to the number and distribution of immune cells within the spleen (Menees et al., 2021) and/or more myeloid-derived suppressor cells in the spleen that infiltrate the CNS (Melero-Jerez et al., 2020). Nevertheless, molecular mechanisms underlying EAE and MS may differ based on the involved neuroanatomical pathways, the disease symptomatology, and the proinflammatory cascade.

The present findings appear to contradict our previous report in male mice where ELS had only a marginal impact on EAE severity (Faraji et al., 2022b). This discrepancy may be explained by sex differences. Sex is a key biological variable that affects vulnerability to stress (Faraji & Metz, 2020) and neuropathological conditions, particularly MS (Doss et al., 2021), and males are less susceptible than females to MS (Voskuhl et al., 2020). Thus, the impact of ELS on the EAE symptoms in female mice here might derive from intrinsic sex differences in response to pathophysiological processes during EAE and stressful experiences (Faraji et al., 2022b; Hoghooghi et al., 2020; Wiedrick et al., 2021). Notably, females and males respond differently to stressful experiences (Faraji et al., 2020a) mainly because of sex differences in their neuroendocrine response by the HPA axis (Oyola & Handa, 2017). In addition, females respond to stress with hyperthermia (Faraji et al., 2022a; Faraji & Metz, 2020), and elevated body temperature can exacerbate symptoms of EAE and MS, including exacerbated postural instability, cognitive dysfunction and impaired vision (Christogianni et al., 2018), also called the Uhthoff phenomenon (Sumowski & Leavitt, 2014).

The present findings show that early life shipment may re-program HPA axis activity with lasting consequences for functional loss and recovery in EAE animals. Early life adversities may disrupt the balance between the function and expression of glucocorticoid receptors (GR) in the brain (de Kloet et al., 2005), induce imbalance between the inhibitory and excitatory actions on the HPA axis (Cotella et al., 2014), and sensitize specific neurocircuits to subsequent acute stressors (Ladd et al., 2005). Both neurohormonal and structural alterations may contribute to the risk for the onset of diseases in later life and reduce psychophysiological resilience that facilitates recovery after adverse stimuli (Cotella et al., 2014). On the other hand, environmental enrichment provides an effective strategy to offset the adverse effects of ELS and build long-lasting resilience (McCreary et al., 2016; McCreary et al., 2019; McCreary & Metz, 2016), however, its effectiveness in the EAE model remains to be shown.

## Conclusion

This study revealed compounding effects of ELS and EAE in adult female mice. The two-hit hypothesis suggests that exposure to ELS may challenge immune or neural systems so that a subsequent insult later in life invokes a potentiated immunological and/or neurological response. It is likely that two hits by stress interact in a synergistic manner to increase the risk or severity of a pathological condition. Here we showed that shipment of laboratory mice in early life re-programs HPA axis function in a way to contribute to earlier and more severe functional loss in an EAE model. The findings suggest that shipment produces transportation stress and potentially skews experimental results and contributes to the replication crisis in the life sciences (Fanelli, 2018). Moreover, the present observations highlight the importance for comprehensive behavioural testing, including non-motor functions such as visual acuity, to enhance the translational value of preclinical animal models of MS. Future studies should investigate differential effects of the two-hit stress model of MS with regard to neuroimmune mediators and mitigation strategies. Thus, psychophysiological stress should be considered a major modifiable risk factor in the onset and progression of MS. The consideration of cumulative effects of lifetime stresses provides a new perspective of the pathogenesis of MS and other autoimmune diseases within a personalized medicine framework.

## Materials and Methods

### Animals

The experimental design is illustrated in Figure 1 (*top row*). Twenty-eight female C57BL/6 [B6] mice, 5–6 weeks old at the beginning of the experiment, were used. The animals were housed in groups of 2-3 per treatment under a 12:12 h light/dark cycle with the light starting at 07:30 h and were provided with water and food *ad libitum*. All experimental procedures were conducted during the light phase of the cycle at the same time of day. To reduce confounding effects of repeated animal handling, the same male experimenters performed all procedures. The experimenters were blinded to the experimental treatments.

Animals were randomly assigned to one of four experimental groups: No Stress (*Control*), Stress (*Control*), No Stress+EAE, and Stress+EAE. All animals were assessed for optokinetic function and exploratory behaviour prior to EAE induction and underwent blood sampling for baseline CORT measurements as the main physiological indicator of HPA axis activation. EAE animals were then immunized in the flanks between postnatal day (P) 48 and 50 (EAE: Day 0) and were assessed for signs of EAE for 26 days. All procedures were carried out in compliance with ARRIVE guidelines and approved by the University of Lethbridge Animal Care Committee in accordance with the standards set out by the Canadian Council for Animal Care (CCAC).

### Early Life Shipment Stress

Mice in stress groups (*n*=17) were shipped from an external breeder (Charles River, Saint-Constant, QC, Canada) to the Canadian Centre for Behavioural Neuroscience (CCBN) in Lethbridge, Alberta at P5-10 as a multidimensional displacement stressor (Poplawski et al., 2023; Poplawski et al., 2020). Briefly, shipped animals were placed into paper crates either the shipping day or the day before, depending upon the time of airplane departure. They were then transported by truck to the airport in a temperature-controlled space. Crates were loaded into the airplane cargo compartment which maintained the ambient climate similar to the passenger cabin (temperature 20°C, pressure 8,000 ft ASL). Flight duration varied from 7 to 9 hours. Upon landing, crates were transported by a passenger car to the destination. It was shown that early life shipment represents a multidimensional stressor resulting in long-term metabolic, neuronal and behavioural changes (Olfe et al., 2010). Upon arrival, animals were immediately placed into home cages and treated in a manner identical to in-house bred animals. No stress groups (*n* =11) were bred in-house at the CCBN.

### EAE Induction and Symptom Monitoring

Mice (No Stress+EAE, *n*=8; Stress+EAE, *n*=11) were weighed prior to EAE induction. EAE induction was performed based on earlier descriptions (Faraji et al., 2022b). Animals’ body weights were also recorded daily to ensure weight loss was within normal bounds (25% of original weight) after EAE induction at P51. Further, using a clinical severity scoring scale (Faraji et al., 2022b), the animals were monitored for 26 days following EAE induction. The clinical scale followed predictable symptom progression from tail paralysis to quadriplegia to death. The summation of disease scores over time (cumulative disease index; CDI) was also calculated from the daily clinical scores (Faraji et al., 2022b). Control (No Stress and Stress) groups received vehicle phosphate-buffered saline (PBS) injections only.

### Visual Acuity and Ocular Reflex Threshold Test

Optic neuritis caused by EAE was assessed through visual acuity and loss of optokinetic head tracking reflexes was measured using the OptoMotry HD ocular reflex threshold system [(Cerebral Mechanics Inc., Medicine Hat, AB, Canada; see also (Prusky et al., 2004)]. The system allows for the measurement of visual acuity and ocular tracking reflexes in awake mice via the optokinetic response to rotating sinusoidal gratings (Douglas et al., 2005). Animals were placed on an elevated platform inside an enclosed chamber with four computer monitors representing the walls of the chamber. A top-down camera was mounted to the top of the chamber to provide a real-time video feed of the animal on the elevated platform. Alternating black and white bars displayed and moved across the monitors, thus creating a virtual rotating cylinder in the chamber. Using the computer terminal for the box, the experimenter monitored the head rotations of the animal, and head rotations in the same direction as the rotating bands indicated that the animal’s visual ability to differentiate the rotating black and white bands was intact. Gradually, the frequency of the bars was increased until the animal could not differentiate the bars and was designated a failed trial. The frequency was then decreased with every failed trial until the animal could differentiate the bars once again. This was performed in the clockwise (CW) and counterclockwise (CCW) direction, randomly alternating directions for each trial, and continued until a frequency threshold (visual acuity) for the animal was found (Alam et al., 2022). It is important to note that the term *acuity* in the present study refers to the maximum spatial frequency that evokes an optomotor response. The animal was then returned to its home cage.

During the course of EAE, optic neuritis was hypothesized to cause severe visual acuity impairments. As such, a maximum of thirty trials was designated based on baseline measurements of the frequency thresholds; typically, a total of 15-20 trials per rotations is required to measure a contrast sensitivity function through each eye. Once this maximum was reached, the last recorded threshold was saved.

### Locomotor Activity and Exploration Task

An AccuScan activity monitoring system (AccuScan Instruments Inc., OH, USA) with clear Plexiglas boxes [length 42 cm, width 42 cm, height 30 cm (Ambeskovic et al., 2019)] was used to assess exploratory activity and affective states in an open field environment. Animals (*n*=10–11/group) were placed individually into the open field and monitored for 5 min at four time points, prior to EAE induction (pre-induction; P42) and on post-induction days 11, 18 and 25. The boxes attached to a computer recorded the activity based on sensor beam breaks. Total travel distance was analyzed using VersaDatTM software (AccuScan Instruments Inc., OH, USA) to measure overall activity. After each animal the apparatus was thoroughly cleaned with 10% Clinicide (Vetoquinol, QC, Canada) to eliminate any odor traces.

### Blood Collection and Spleen Weights

Baseline blood samples were collected 7 days prior to EAE induction at P43 and during the peak of symptoms at 19 days post-induction (day19). Briefly, in manually restrained mice, a small puncture was made in the sub-mandibular vein using a lancet, and approximately 0.1 ml of blood was collected with a micro-tube. All samples were collected in the morning hours between 09:00 and 11:00 h. Animals were returned to their home cage and allowed to recover for 4 days. Post-recovery blood samples were collected immediately prior to euthanization (day27) through a cardiac puncture using a heparin coated syringe. Animals were euthanized through an intracardiac injection of Euthansol (pentobarbital, 150 mg/kg; CDMV Inc., Saint-Hyacinthe, QC, Canada). Plasma was obtained by centrifugation at 5,000 rpm for 10 min and stored at −80°C. Spleen was removed and weighed on an electronic scale to determine the possible impact of ELS and EAE on the organ central to endocrine and immune regulation (Faraji et al., 2022a).

### Data Analysis

In all statistical analyses (SPSS 16.0, SPSS Inc., USA), a *p*-value of <0.05 (two-tailed) was chosen as the significance level, and results are presented as mean ± standard error. EAE disease scores, optokinetic responses and CORT levels were often not normally distributed, and Levene’s test did not confirm homogeneity of variance in the spleen weights. Therefore, Kruskal-Wallis H test, a rank-based nonparametric test for analysis of variance was used to detect differences between means of groups across multiple test trials. The Mann-Whitney *U* test was also used to compare means of the two groups (No Stress+EAE and Stress+EAE) for a single dependent variable (EAE disease score and CDI) with Bonferroni correction for multiple comparisons when necessary. Correlation between variables (CDI, distance traveled, frequency threshold, CORT, spleen weight, etc.) was analyzed by Spearman’s rank correlation. Because Spearman’s rank correlation is more resistant to outliers, it was also used when one variable was normally distributed. Also, *Repeated measures (R-M)* and *one-way (O-W)* ANOVA were employed to analyse distance traveled in the open field task and frequency threshold. *Post-hoc* test (Tukey HSD) was used to adjust for multiple comparisons. In all cases, means of values were compared. To evaluate the magnitudes of the effects, effect sizes (η2 for ANOVA) were calculated. Values of η2 = 0.14, 0.06, and 0.01 were considered for large, medium, and small effects, respectively.

## Author contributions

DB and GASM designed the study. DB and JF performed the experiments and analyzed the data. JF wrote the paper. DB, VWY and GASM edited the paper. VWY and GASM acquired the funding and provided resources.

## Acknowledgments

The authors acknowledge support by the Alberta Innovates-Health Solutions Collaborative Research and Innovation Opportunities (CRIO) program (VWY, GM), Natural Sciences and Engineering Research Council (NSERC) of Canada Discovery Grants #5628 and #31 (GM), and Canadian Institutes of Health Research (CIHR) Project Scheme #363195 (GM).

